# RelCoVax®, a two antigen subunit protein vaccine candidate against SARS-CoV-2 induces strong immune responses in mice

**DOI:** 10.1101/2022.01.07.475330

**Authors:** Abhishek Phatarphekar, G. E. C. Vidyadhar Reddy, Abhiram Gokhale, Gopala Karanam, Pushpa Kuchroo, Ketaki Shinde, Girish Masand, Shyam Pagare, Nilesh Khadpe, Sangita S. Pai, Vijita Vijayan, R. L. Ramnath, K. Pratap Reddy, Praveen Rao, S. Harinarayana Rao, Venkata Ramana

## Abstract

The COVID-19 pandemic has spurred an unprecedented movement to develop safe and effective vaccines against the SARS-CoV-2 virus to immunize the global population. The first set of vaccine candidates that received emergency use authorization targeted the spike (S) glycoprotein of the SARS-CoV-2 virus that enables virus entry into cells via the receptor binding domain (RBD). Recently, multiple variants of SARS-CoV-2 have emerged with mutations in S protein and the ability to evade neutralizing antibodies in vaccinated individuals. We have developed a dual RBD and nucleocapsid (N) subunit protein vaccine candidate named RelCoVax® through heterologous expression in mammalian cells (RBD) and *E. coli* (N). The RelCoVax® formulation containing a combination of aluminum hydroxide (alum) and a synthetic CpG oligonucleotide as adjuvants elicited high antibody titers against RBD and N proteins in mice after a prime and boost dose regimen administered 2 weeks apart. The vaccine also stimulated cellular immune responses with a potential Th1 bias as evidenced by increased IFN-γ release by splenocytes from immunized mice upon antigen exposure particularly N protein. Finally, the serum of mice immunized with RelCoVax® demonstrated the ability to neutralize two different SARS-CoV-2 viral strains *in vitro* including the Delta strain that has become dominant in many regions of the world and can evade vaccine induced neutralizing antibodies. These results warrant further evaluation of RelCoVax® through advanced studies and contribute towards enhancing our understanding of multicomponent subunit vaccine candidates against SARS-CoV-2.

## 1. Introduction

In the past two decades, coronaviruses (CoV) have emerged as a serious threat to humans with 3 major outbreaks being reported starting with severe acute respiratory syndrome (SARS) in late 2002, the Middle Eastern respiratory syndrome (MERS) in 2012 and coronavirus disease (COVID-19) in December 2019 ^[1–3]^. Of these, COVID-19 pandemic caused by the SARS-CoV-2 virus has been the most devastating with greater than 250 million infections and over 5 million deaths reported globally within 2 years of the outbreak first being reported in Wuhan, China ^[4]^. A massive global collaborative effort was needed to develop safe and effective vaccines to protect the world population from the rampaging SARS-CoV-2 virus ^[5]^. Multiple vaccine candidates were developed with unprecedented agility using novel platform technologies such as mRNA and carrier viral vector based vaccines as well as traditional platforms such as the inactivated whole virus and protein subunit vaccines ^[6]^.

Most of the approved vaccines as well as vaccine candidates in development, elicit an immune response against the spike (S) surface glycoprotein of SARS-CoV-2 that recognizes the angiotensin converting enzyme 2 (ACE2) receptor on the target cell via the receptor binding domain (RBD) and enables viral entry ^[7,8]^. Hence antibodies binding to RBD have the potential to block the RBD-ACE2 interaction and neutralize SARS-CoV-2 viral entry. Indeed, it has been observed that sera from rabbits immunized with RBD had a higher neutralizing antibody titer compared to sera from rabbits immunized with full S protein or S1 domain of S protein ^[9]^ and higher anti-RBD antibody titer correlated with higher neutralizing antibody titer in patient sera ^[10]^.

Within the past year, SARS-CoV-2 variants with mutations in S protein especially in RBD have emerged with the potential to evade neutralizing antibodies in vaccinated individuals as highlighted in certain *in vitro* neutralization studies ^[11]^. One particular SARS-CoV-2 variant known as the Delta variant (B.1.617.2) has been of particular concern spreading across multiple geographies and demonstrating the potential to escape neutralization by antibodies elicited through immunization with multiple vaccines targeting the S protein ^[12,13]^. Hence there is a need for next generation vaccines that are able to protect against multiple variants and encompass additional antigens that have a lower potential for mutation thereby providing broader protection. One such candidate is the nucleocapsid (N), a structural protein required for packaging the viral RNA [14]. The amino acid sequence of N protein is well conserved with 90% homology between SARS-CoV and SARS-CoV-2 ^[15]^. Convalescent patient serum samples have shown significant N reactive T-cells indicating the potential for stimulating strong cell mediated immunity and long term immunity ^[16]^.

Often, protein subunit vaccine candidates are weak immunogens and require formulation with adjuvants to boost immune response. Aluminum compounds (aluminum hydroxide or alum and aluminum phosphate) are commonly used as adjuvants and boost humoral immune responses but do not stimulate cell mediated immunity adequately ^[17,18]^. Toll like receptor 9 (TLR9) agonists such as CpG oligonucleotides are known to stimulate a Th1 biased T-cell response ^[19]^ and can be used in combination with aluminum based adjuvants to stimulate both humoral and cell mediated immune responses.

We developed RelCoVax®, a RBD and N protein two antigen SARS-CoV-2 vaccine candidate. The vaccine was designed to provide broader protection against SARS-CoV-2 variants and in this study we demonstrate that the vaccine is strongly immunogenic in mice and produces antibodies that can neutralize two strains of SARS-CoV-2 including the Delta variant.

## 2. Materials and Methods

### 2.1. Expression of proteins

The published sequence of the SARS-CoV-2 virus Wuhan strain (NCBI Accession MN908947.3) ^[20]^ was used as a reference for protein sequences of S (residues 1-1273) and N (residues 1-419) proteins. DNA sequences for S and N proteins were codon optimized for expression in *Cricetulus griseus* (Chinese hamster) ovary or CHO cells and *E. coli* respectively and synthesized by GeneArt (Thermofisher Scientific). The DNA sequence for RBD (Arg 319 to Phe 541) was amplified from the synthetic S gene and cloned into the *EcoRI* and *HindIII* digested pXC17.4 CHO expression vector with a secretion signal sequence using NEBuilder HiFi DNA assembly kit (New England Biolabs, E5520). Similarly, the DNA sequence for the synthetic N gene (Met1 to Ala 419) was amplified and cloned into the *NdeI* and *XhoI* digested pET24a(+) vector (Novagen).

The pXC17.4-RBD plasmid was transfected into CHOK1SV GS-KO cells (Lonza) by electroporation and stably transfected clones were selected by addition of methionine sulphoxamine (Sigma Aldrich) to the CD-CHO media (Gibco). The supernatants of individual clones were screened for expression of RBD qualitatively by ELISA using the GENLISA™ Human SARS-CoV-2 (COVID-19) spike protein S1 antigen ELISA kit. (Krishgen Biosystems, KBVH015-10). The highest RBD expressing clone was transferred to the bioreactor for upstream process development.

The pET24a(+)-N plasmid was transformed into chemically competent BL21DE3 pLysS *E. coli* cells (Novagen) and the transformants were isolated on kanamycin containing LB plates. Single colonies were inoculated and protein expression was induced using isopropyl-β-D-thiogalactopyranoside (IPTG, Sigma Aldrich). Cell lysates were screened for protein expression using SDS-PAGE followed by Coomassie blue staining. A single N expressing clone was transferred to the fermenter for upstream process development.

### 2.2. Purification of RBD

SARS-CoV-2 RBD was expressed during fermentation by the stably transfected CHOK1SV-GS KO cell line and secreted extracellularly. The cell free supernatant derived during harvest of the fermented broth was processed using Blue Sepharose™ 6 Fast Flow column (GE Healthcare, Cat# 17-0948-01) in flow-through mode followed by a hydrophobic interaction chromatography (HIC) purification step. The protein was then dia-filtered into a low salt buffer and further processed by anion exchange chromatography. The eluted protein was dia-filtered into the storage buffer, processed for a final endotoxin removal step, sterile filtered and stored at −20 (±5) °C.

### 2.3. Purification of N protein

SARS-CoV-2 Nucleocapsid was expressed as an intracellular protein in *E.coli.* The cells were lysed by homogenization and the lysate was centrifuged to obtain the cell free supernatant containing the protein of interest. The supernatant was subjected to depth filtration and processed by anion exchange chromatography in flow through mode. After dia-filtration, the protein was further processed by cation exchange chromatography and the fractions were pooled after analysis of purity by reverse phase (RP) HPLC. The pooled fractions were buffer exchanged into the storage buffer, processed for a final endotoxin removal step, sterile filtered and stored at −70 (±5) °C.

### 2.4. General materials

In western blots, RBD was detected using SARS-CoV-2 Spike RBD antibody (Prosci # 9087) as primary antibody and goat anti-rabbit alkaline phosphatase (Jackson Immunoresearch # 111-055-003) as secondary antibody. N protein was detected using SARS-CoV-2 nucleocapsid antibody (Prosci # 35-579) as primary antibody and goat anti-mouse IgG whole molecule (Sigma # A4416) as secondary antibody.

Aluminum hydroxide gel (Alhydrogel®) or alum adjuvant was purchased from Croda, Denmark (Cat # AJV3012) and the CpG deoxy oligonucleotide with a phosphorothioate backbone was custom synthesized by Synbio Technologies, NJ, USA.

### 2.5. RBD and ACE2 interaction studies

To evaluate the binding of RBD to ACE2 receptor, SPR based assays were performed using the Biacore™ T200 (GE Healthcare) system and the kinetic rate constants (k_a_ and k_d_) and the affinity constant (K_D_) were determined for 3 individual batches of purified RBD protein. Human ACE2 with Fc tag (Acro Biosystems, AC2-H5257) was captured on the CM5 chip using a running buffer comprising of 10 mM HEPES pH 7.2, 150 mM NaCl and 0.05% Tween-20. The RBD samples were serially diluted from 50 nM to 3.125 nM in running buffer and passed over the chip surface. Single-cycle kinetics was performed using 1:1 kinetic binding model and the sensograms were evaluated using the BIA evaluation software.

### 2.6. Animal ethics, handling and treatment

Adult male and female BALB/c mice (inbred) were housed at Laboratory Animal Research Services, Reliance Life Sciences, Navi Mumbai. Animals were certified by the veterinarian to be free of any illness. Animals were acclimatized for at least 5 days before initiation of the study. The temperature was maintained at 22 ± 3°C, relative humidity at 50 ± 20% with a light / dark cycle of 12 hours each. Mice had free access to *ad libitum* standard rodent diet and water. Studies were performed in accordance with the recommendations of the Committee for the Purpose of Control, and Supervision of Experiments on Animals (CPCSEA) guidelines published in The Gazette of India, 2018 (Compendium of CPCSEA, 2018). The experimental protocols were approved by institutional animal ethical committee (IAEC) and studies were performed as per the standard operating procedures of animal house facility.

A total of 4 different studies were performed and animals in the studies were initially randomized into different groups based on body weight. Body weight variation was within ± 20% of the mean body weight for any sex. The route of injection was intramuscular and dose volume was either 100 μL or 200 μL (50 μL or 100 μL in each hind leg) for both primary and booster doses in all studies. In all vaccine formulations, 500 μg of alum and 200 μg of CpG oligonucleotide were added as adjuvants per dose volume. Blood was collected in inert gel tubes using mild anesthesia [0.2% (v/v) diethyl ether] via open drop method, allowed to settle for at least 30 minutes at room temperature and centrifuged to collect the serum. Animals were observed for mortality and signs of morbidity at least twice a day and were monitored carefully for any signs of illness or reaction towards treatment once a day after dose administration till the completion of the experimental period. The injection sites of all animals were observed for erythema and oedema once daily until study completion. All animals were checked for detailed clinical observations, ophthalmological examinations, body weight and feed consumption before treatment and then weekly till the completion of study.

### 2.7. Estimation of anti-RBD and anti-N antibodies in mouse sera by ELISA

Anti-RBD and anti-N antibodies were estimated in the sera of immunized mice by using the end-point titer ELISA method. Briefly, 96 well ELISA plates (Nunc Maxisorp, 44-2404-21) were coated with 300 ng of respective antigen in phosphate buffered saline (PBS) and incubated overnight at 4 °C. Plates were washed once with PBS containing 0.1% Tween 20 (PBS-T) and blocked with 3% bovine serum albumin (BSA, Sigma Aldrich) in PBS-T for 1 hr at room temperature. After blocking, the 3% BSA solution was discarded and mouse serum diluted with 0.1% BSA in PBS-T was added to the top row of each plate, serially diluted further down each column and incubated at 37 °C for 2 hrs. Plates were washed thrice with PBS-T followed by addition of peroxidase conjugated goat anti-mouse IgG diluted 15,000 fold and 20,000 fold (Sigma Aldrich, A4416) to RBD and N protein coated plates respectively. After incubation at 37 °C for 1 hr, plates were washed 3 times with PBS-T and TMB substrate (Denovo Biolabs) was added for development of color. Reaction was stopped after 30 mins by addition of 1N sulphuric acid. Optical density (OD) was measured at 450 nm and 620 nm using Powerwave XS ELISA Plate reader (Biotek Instruments). For calculations, OD (450 nm - 620 nm) value was used for each well.

### 2.8. SARS-CoV-2 neutralization assays

The plaque reduction neutralization tests (PRNT50) were performed at National Immunogenicity and Biologics Evaluation Centre (NIBEC), Bharati Vidypeeth, Pune - 411043, Maharashtra, India as per established standard operating procedures. All steps involving live virus handling were performed in a BSL-3 facility. Immunized mice sera were tested against SARS-CoV-2-IND/8004/2020 (B.1.1.306) and SARS-CoV-2-IND/2095/2021 (B.1.617.2, Delta) strains. For this assay, serum samples from 4 mice were pooled together in serial order to generate 3 individual pools of sera for each test group and placebo group consisting of 12 mice each. Briefly, Vero (CCL-81) cells were seeded in 24 well plates (10^5^ cells per well in MEM + 10% FBS) and grown to about 90% confluency prior to starting the assay. Test serum samples along with controls were diluted in MEM + 2% FBS and heat inactivated at 56 °C for 30 mins. Serial dilution of serum samples was performed in a 96 well plate. The viral stocks were diluted in media, added to the wells containing serum dilutions (1:1 ratio at 60 to 100 pfu viral titer per well) and incubated for 1 hr at 37 °C and 5% CO2. After incubation, these were transferred in duplicate to the 24 well plate containing Vero cells after removal of growth media and incubated for 1 hr at 37°C and 5% CO2 with mixing every 15 mins for viral adsorption. The final viral titer per well was between 30-50 pfu. One mL of overlay media containing carboxy methyl cellulose was added to the wells and the plates were incubated for 5 days at 37 °C and 5% CO2. Wells were washed once with PBS and fixed with formaldehyde solution. After 3 washes, staining was performed by addition of 1% crystal violet solution to the wells. Plates were washed thrice with distilled water, dried and the plaques were counted. The mean number of plaques from the virus only control wells was used to determine the cutoff for desired level of infection reduction. PRNT50 was reported as the highest dilution of test serum for which virus infectivity was reduced by 50%. A standard logical regression model was followed for determination of PRNT50 and cut-off values for determination of valid data points were calculated using the NIH LID Statistical web tool (https://bioinformatics.niaid.nih.gov/plaquereduction/).

### 2.9. ELISpot assay

To determine the frequency of IFN-γ producing cells by the vaccine, we used antigen-specific IFN-γ ELISpot kits (MABTECH, 3321-4HPT-10) as per the manufacturer’s instructions. Briefly, spleens from mice were transferred to complete RPMI (RPMI + 10% FBS). The spleens were then individually crushed using sterile frosted slides in petri dishes containing complete RPMI, passed through a 40 μm cell strainer (Becton Dickinson), collected in sterile centrifuge tubes and pelleted. The cells were resuspended in ACK lysing buffer (Gibco, A1049201) and incubated for 5 mins at room temperature followed by addition of 10% FBS in PBS. The cells were further washed 3 times with 10% FBS in PBS, resuspended in complete RPMI, counted and plated in IFN-γ mAb (AN18) coated plates at 1 million cells per well and 0.5 million cells per well in duplicate. The cells were stimulated by RBD, N protein and RBD plus N protein together each at a concentration of 10 μg/mL along with ConA (10 μg/mL) as positive control and only media as negative control for each group. The plates were incubated at 37 °C for 40 to 42 hrs. Subsequently, the plates were washed and incubated with a biotinylated detection antibody, followed by Streptavidin-HRP. The plates were developed with the TMB substrate as per the manufacturer’s instructions until distinct spots emerged. The plates were imaged using ImmunoSpot reader (Cellular Technologies Ltd) and number of spots per well were counted using Biospot 5.0 software.

### 2.10. Data Analysis

All data was analyzed and graphically represented by using Graphpad prism (9.2.0). Statistical significance was determined using unpaired two-tailed T-test and p-values < 0.05 were considered statistically significant (* p-value < 0.05, ** p-value < 0.005, *** p-value < 0.0005, NS - not significant). All preliminary data was tabulated and basic calculations were performed using Microsoft Excel.

## 3. Results

### 3.1. Characterization of RBD and N proteins

We expressed a part of the S protein extending from residue 319 to 541 (223 amino acids) encompassing the RBD and the full length N protein (419 amino acids) (Fig. 1A). Both RBD and N protein were purified to homogeneity through multistep purification processes. Under nonreducing conditions, the RBD protein band was observed between 24 and 31 KDa (Fig. 1B) indicating monomeric form of the protein (MW = 25 KDa). The N protein sequence lacks cysteine residues and was observed at the expected size below 52 KDa (MW = 45.6 KDa). To determine whether the expressed RBD protein has folded correctly and is able to bind to its target, its interaction with the ACE2 receptor was probed. We observed that RBD bound to ACE2 with high affinity in the low nM range (Table 1) well within the reported K_D_ range of 5 nM to 70 nM for RBD-ACE2 interaction ^[7,21] [22]^.

**Fig. 1.**
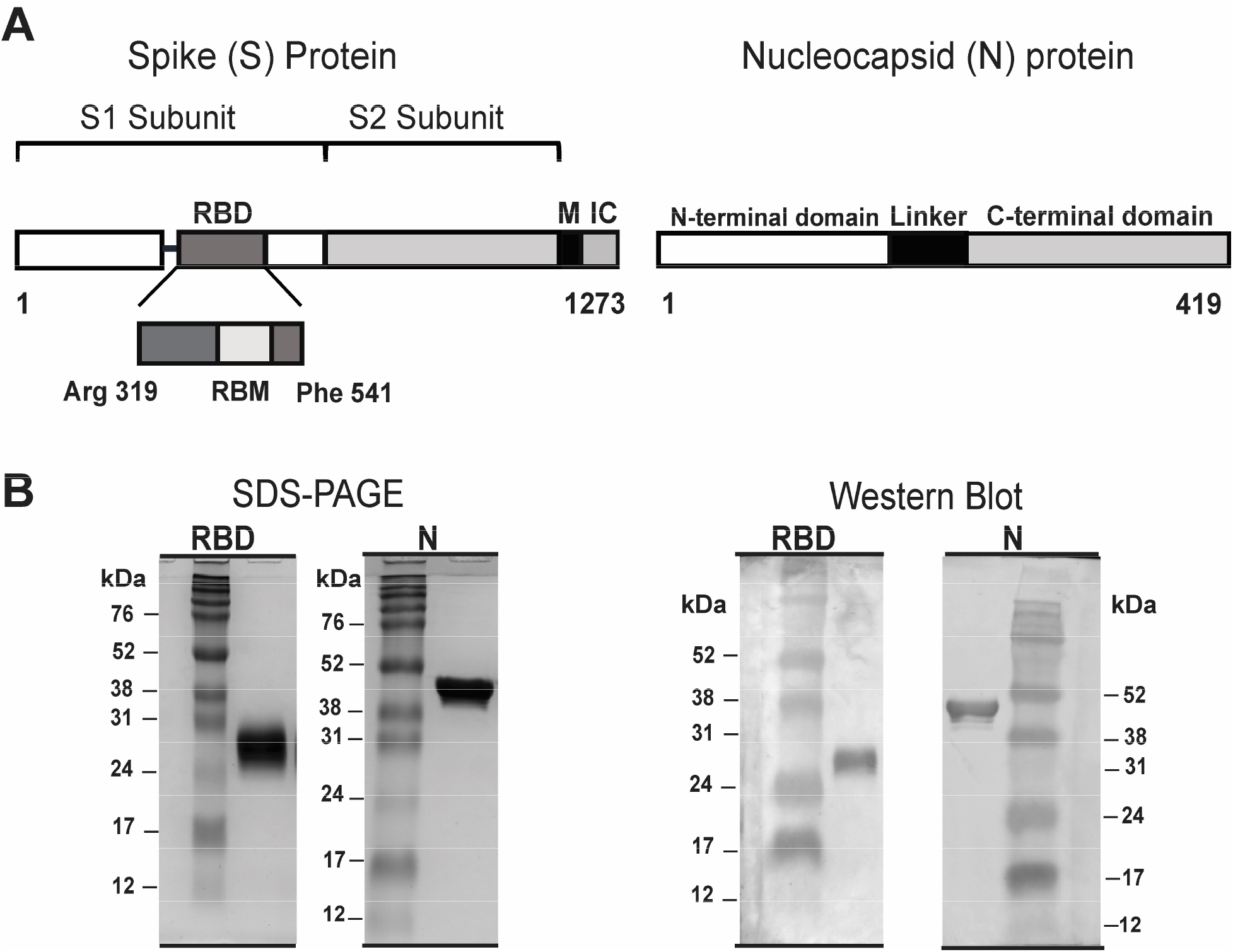
Expression of RBD and N proteins. (A) Schematic representations of S and N proteins of SARS-CoV-2. Receptor binding domain or RBD of S protein as expressed extends from Arg 319 to Phe 541 and includes the receptor binding motif (RBM) ^[7]^. M is the transmembrane domain and IC is the intracellular domain of S protein. For N protein, full length protein is expressed. (B) Representative SDS-PAGE and western blot images for purified RBD and N proteins. Molecular weights the for reference ladder are indicated in kDa (kilodaltons).

**Table 1.** Affinity constants for interaction of RBD and ACE2 as measured by SPR. Three independent batches of purified RBD were evaluated for ACE2 binding.

| **RBD** | **$k_a$ ($M^{-1}s^{-1}$)** | **$k_d$ ($s^{-1}$)** | **$K_D$ (M)** | **$K_D$ (nM)** |
| --- | --- | --- | --- | --- |
| Batch 1 | $6.24 \times 10^5$ | $1.14 \times 10^{-2}$ | $1.83 \times 10^{-8}$ | 18.3 |
| Batch 2 | $7.63 \times 10^5$ | $1.15 \times 10^{-2}$ | $1.51 \times 10^{-8}$ | 15.1 |
| Batch 3 | $6.19 \times 10^5$ | $1.11 \times 10^{-2}$ | $1.79 \times 10^{-8}$ | 17.9 |
| Mean | $6.69 \times 10^5$ | $1.13 \times 10^{-2}$ | $1.71 \times 10^{-8}$ | 17.1 |

### 3.2. Clinical Observations

No mortality or morbidity was observed in any of the animals due to treatment in all studies. One animal died on day 21 during blood collection in the 1 μg antigen test group in the dose comparison study. Slight erythema and oedema was observed at the site of injection and it recovered by end of the experimental period. No significant differences were observed in body weight and feed consumption of animals in different groups of each study. No treatment related changes were observed in detailed clinical and ophthalmological examinations in any of the studies.

### 3.3. Humoral immunity induced by RelCoVax® in BALB/c mice

Initially we studied the effect of adjuvants alum and CpG oligonucleotide on anti-RBD and antiN antibody titers in mice sera. The mice were immunized and serum collected as per schematic in Fig. 2A. With no adjuvants (G1), RBD was poorly immunogenic and did not produce significant antibody titers even after 2 doses compared to the placebo group (P) (Fig. 2B). When alum was included as an adjuvant (G2 and G4), a strong stimulation of anti-RBD antibody titer was observed compared to other groups. The highest anti-RBD antibody geometric mean titer (GMT) was observed for G4 when both alum and CpG oligonucleotide were added as adjuvants. The difference between the anti-RBD titer for G2 and G4 was statistically significant after 1^st^ dose but not after the 2^nd^ dose. Unlike RBD, mice in group G1 displayed significantly higher anti-N antibody titer compared to mice in group P indicating that N protein is immunogenic even in absence of adjuvants (Fig. 2C). Similar to RBD, mice in the groups G2 and G4 displayed a sharp increase in the anti-N antibody titers compared to other groups indicating strong stimulation with alum. Overall, G4 mice displayed the highest anti-N antibody titers with strong statistical difference compared to G2 mice after 1^st^ dose but not after 2^nd^ dose.

**Fig. 2.**
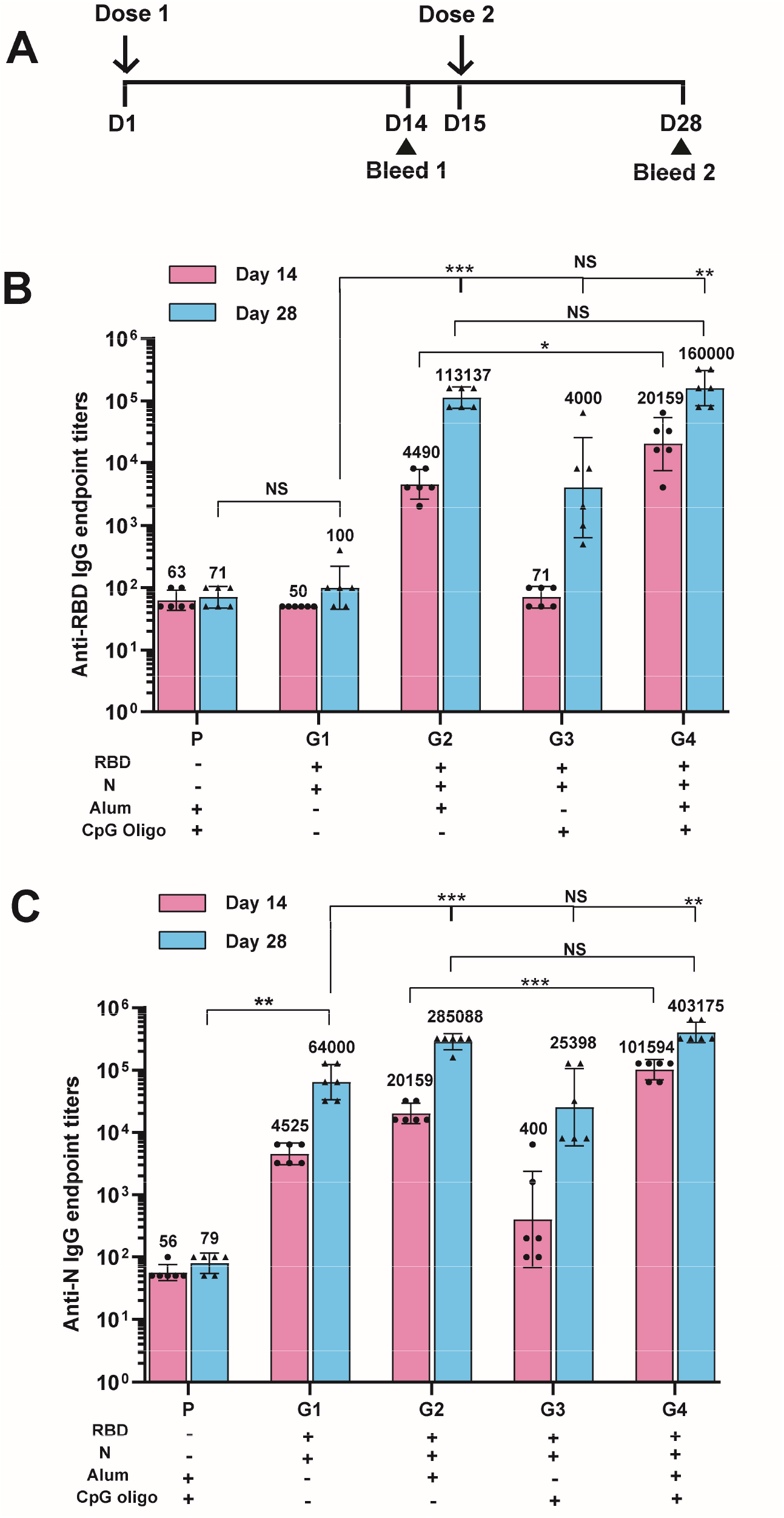
Effect of adjuvants on immunogenicity of RBD and N proteins. (A) Dosing schedule for mice (N=6 per group) used in this study. Amount of each antigen is 10 μg formulated with or without adjuvants in a total volume of 100 μL per dose. Two doses for each formulation were administered 14 days apart (day 1 and day 15) and serum samples were collected 13 days after each dose. (B) Anti-RBD IgG and (C) anti-N IgG end point titers for each mouse group. P indicates placebo group while G1 to G4 represent test groups. (+) and (-) signs indicates the presence and absence of each component in the formulation. The bars and the numbers indicate the geometric mean titer (GMT) for each group and error bars represent the 95% confidence interval (CI) for that group.

We also evaluated the anti-RBD and anti-N antibody titers in mice sera with a 10-fold difference in dose of each antigen (1 μg vs 10 μg) and tracked the titers weekly as per schematic in Fig. 3A. Significant increase in titer of anti-RBD and anti-N antibodies compared to placebo could be detected in serum collected on day 14 nearly 2 weeks after the priming dose (Figs. 3B and 3C). Peak anti-RBD and anti-N antibody levels were measured in serum samples collected on day 21 nearly a week after the booster dose. Overall, 10 μg of antigen per dose produced higher antibody titers than 1 μg of antigen for both RBD and N proteins. For anti-RBD antibody titers, statistical significance was observed between the 2 dosing regimens only on day 14 (after 1^st^ dose) (Fig. 3B). For anti-N antibody titers, statistical significance was observed between the 2 dosing regimens on day 14 as well as on day 28 due to an approximately 3-fold decrease in anti-N GMT between day 21 and day 28 for the 1 μg dose group (Fig. 3C).

**Fig. 3.**
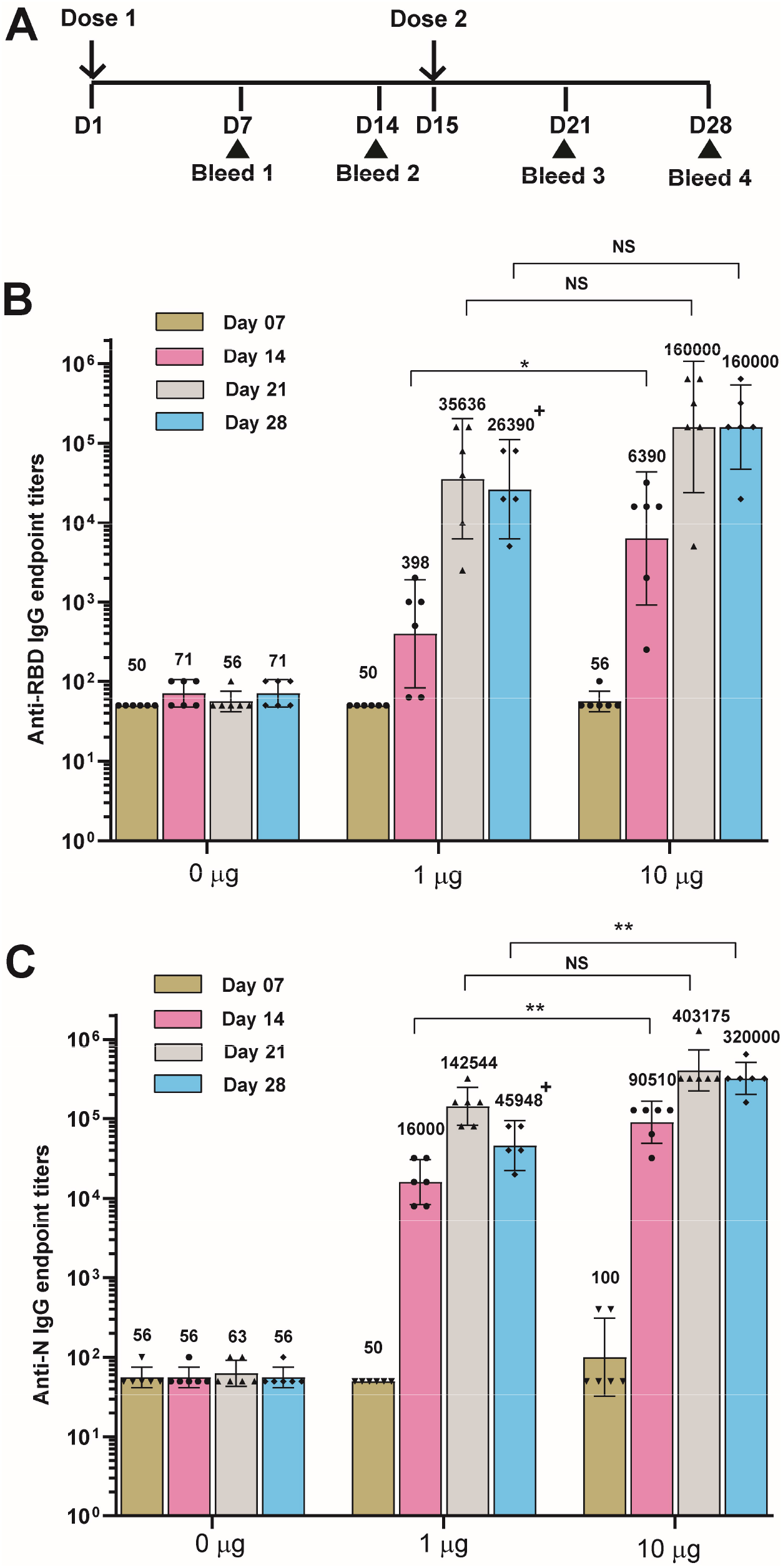
Comparison of 1 μg and 10 μg antigen doses. (A) Dosing schedule for mice (N=6 per group) to compare 1 μg and 10 μg doses of each antigen (no antigen is placebo group). All formulations contain alum and CpG oligonucleotide as adjuvants in a total volume of 100 μL per dose. Two doses for each formulation were administered 14 days apart (day 1 and day 15) and serum samples were collected 6 days and 13 days after each dose. (B) Anti-RBD IgG and (C) anti-N IgG end point titers for each mouse group. The bars and the numbers indicate the GMT for each group and error bars represent the 95% CI for that group. (+) indicates N=5 since one animal in the 1 μg dose group died during blood collection on day 21 (no serum sample on day 28 for that mouse).

We optimized the formulation of our vaccine candidate to include a preservative and had to increase the minimum formulation volume to 200 μL per dose while maintaining the same amount of each antigen (10 μg) and adjuvants per dose. We also formulated RelCoVax® in a volume of 500 μL per dose envisaged for humans. In subsequent studies, the anti-RBD and anti-N antibody response induced by both these formulations was studied following 2 doses each per mouse (Fig. 4A). For the 500 μL formulation only 40% of the volume could be injected into each mouse (maximum permissible volume is 200 μL per animal) per dose implying that each animal received only 40% of each antigen (4 μg) and adjuvant compared to the 200 μL formulation (10 μg). We observed that the anti-RBD as well as anti-N antibody titers were comparable between the 200 μL formulation and the 500 μL formulation of the vaccine (Fig. 4B). Interestingly, anti-RBD antibody titers for both these formulations were elevated by about 2 to 3 fold (normalized to placebo) compared to the 100 μL formulation volume used in earlier studies (Figs. 2B and 3B). The difference was less pronounced for anti-N antibody titers (1.5 to 2 fold normalized to placebo) (Figs. 2C and 3C).

**Fig. 4.**
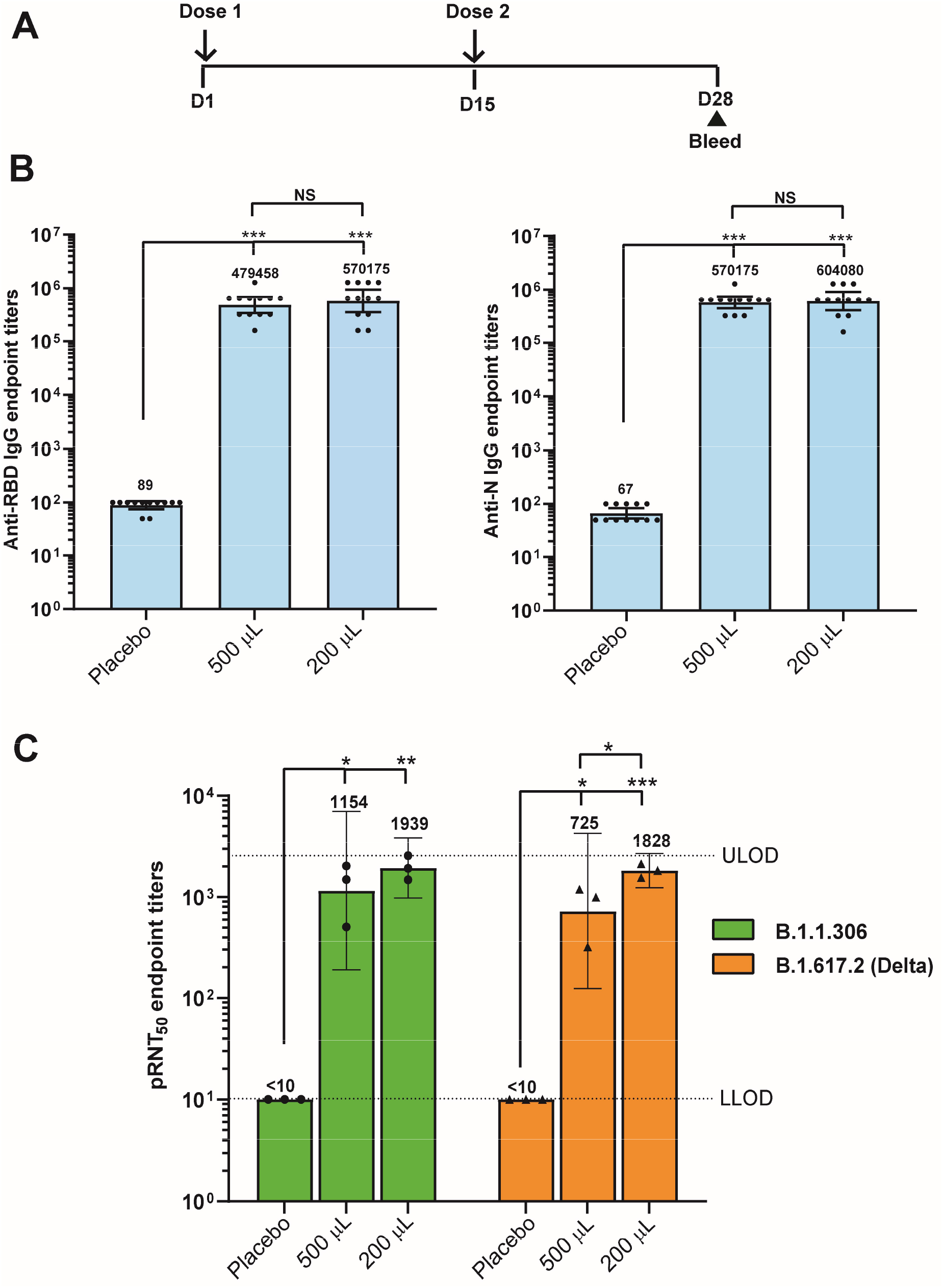
Immunogenicity of 200 μL and 500 μL formulations of RelCoVax®. (A) Dosing schedule for mice (N=12 per group) receiving two different formulations (200 μL per dose volume and 500 μL per dose volume) of vaccine candidate with no antigen placebo group as control (200 μL dose volume). Two doses for each formulation were administered 14 days apart (day 1 and day 15) and serum samples were collected on day 28 (13 days after dose 2). The two formulations contained 10 μg of each antigen along with alum and CpG oligonucleotide as adjuvants. Each mouse was administered 200 μL from either formulation per dose. (B) Anti-RBD IgG and anti-N IgG end point titers for each mouse group. The bars and the numbers indicate the GMT for each group and error bars represent the 95% CI for that group. (C) SARS-CoV-2 neutralizing endpoint titers (pRNT50) for 3 serum pools (4 mice per pool). The serum pools were prepared from the same serum samples tested in Fig. 4B. The bars and the numbers indicate the GMT for each group and error bars represent the 95% CI for that group. LLOD – lower limit of dilution, ULOD – upper limit of dilution.

We further tested the above serum pools (3 pools of 4 mice each per dose group) for *in vitro* neutralization against both the B.1.1.306 and the Delta strains of SARS-CoV-2 in a PRNT 50 assay. Serum dilutions for PRNT50 ranged from 10 fold to 2560 fold hence these were the limits of quantification for the titer values. Relative to placebo, significantly stronger virus neutralization was observed in test serum pools indicating presence of neutralizing antibodies against both strains of the virus (Fig. 4C). Immunization with the 200 μL formulation produced higher overall neutralizing antibody titer (statistical significance was observed between the 200 μL formulation and the 500 μL formulation against the Delta strain).

### 3.4. Cell mediated immunity induced by RelCoVax® in BALB/c mice

Cellular responses to RBD and N proteins in immunized animals as well as control group dosed as per schematic in Fig. 5A were determined by stimulating splenocytes with both antigens individually as well as in combination. Media only was used as negative control and Con A was used as a positive control (data not shown). We observed higher overall IFN-γ positive cells per million splenocytes in test mice compared to placebo mice across all treatment groups (Fig. 5B). In the test group, significantly higher stimulation was seen in N protein and RBD plus N protein treated group compared to media treated group (p-values of 0.0097 and 0.0098 respectively). We did not see much IFN-γ response in media treated and RBD treated placebo mice splenocytes but N protein and RBD plus N protein treatment resulted in high IFN-γ response even in placebo group. However, the test mice splenocytes showed significantly higher IFN-γ response to N protein and RBD plus N protein compared to placebo group (p-values of 0.015 and 0.0023 respectively).

**Fig. 5.**
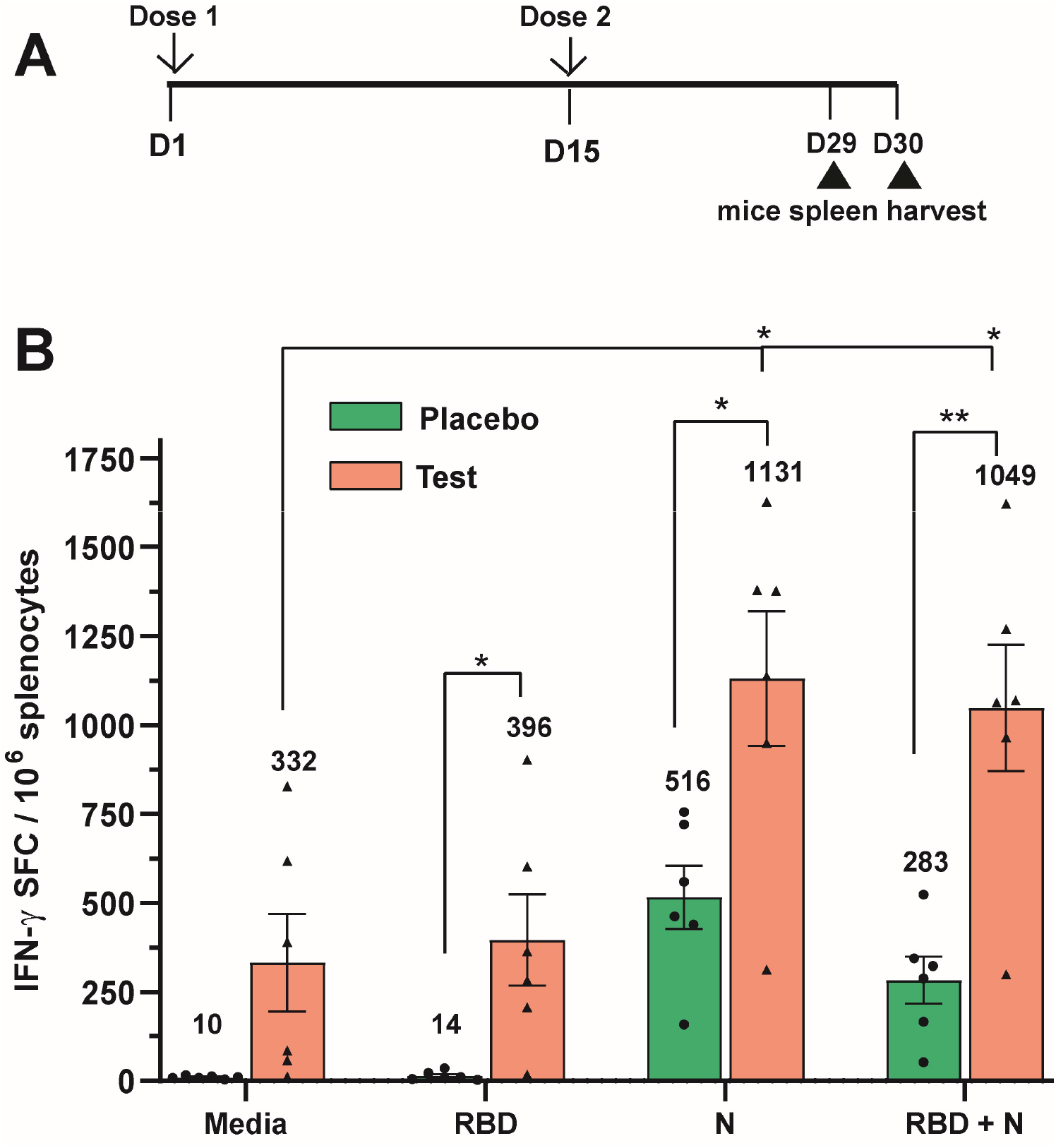
Assessment of cellular immune responses. (A) Dosing schedule for mice (N=6 per group) receiving the vaccine formulation containing 10 μg of each antigen with alum and CpG oligonucleotide added as adjuvants in 200 μL dose volume. No antigen control (200 μL) was used as placebo. Two doses were administered 14 days apart (day 1 and day 15) and 3 mice from each group were sacrificed day 29 and day 30 respectively. Splenocytes were harvested after sacrifice for ELISpot assay. (B) Cellular immune responses as measured by IFN-γ ELISpot of splenocytes from immunized mice when stimulated with different antigens. The bars and the numbers indicate the mean number of IFN-γ spot forming cells (SFCs) per million splenocytes for each group and error bars indicate the standard error of mean (SEM).

## 4 Discussion

With the COVID-19 pandemic caused by the SARS-CoV-2 virus continuing to spread across the world including in some countries where a substantial population has been vaccinated, there is a need for development of vaccines targeting multiple viral antigens. In this paper we have described the development of RelCoVax®, a two antigen adjuvanted subunit vaccine candidate comprising of the RBD and N proteins of SARS-CoV-2 virus and its immunogenicity in mice. Vaccine treatment did not affect the body weight or feed and water consumption and no treatment related clinical signs of toxicity were observed during experimental period in any of the animals barring transient local injection site reactions indicating that the vaccine was generally safe for the treated animals.

Previous studies with the SARS-CoV virus have indicated that RBD produced through heterologous expression and adjuvanted with alum can neutralize viral entry into cells ^[23]^. In addition to alum, we also included a CpG oligonucleotide, an adjuvant known to stimulate a Th1 biased immune response with stronger stimulation of cell mediated immunity. This combination of adjuvants has also been tested with SARS-CoV-2 subunit vaccine candidates based on the full length S protein ^[24,25]^. Our vaccine candidate produced a robust antibody response to both RBD and N antigens in immunized mice with endpoint titers well over 1000 fold higher compared to placebo treated mice after a 2 dose regimen. Elevated anti-RBD and anti-N antibody levels could be detected nearly 2 weeks after first dose and the second dose led to a strong increase in titer to peak levels within a week after administration. A dose of 10 μg of each antigen produced higher titers than 1 μg of each antigen. However, we did observe a dose sparing effect when formulation volumes were readjusted to include a preservative. Here, a dose of 40% of each antigen and adjuvant produced nearly the same levels of anti-RBD and anti-N antibody titers (200 μL Vs 500 μL formulation).

There is a strong rationale for increasing the coverage of SARS-CoV-2 antigens beyond the S protein with new mutations being identified in the S protein including the RBD in emerging variants, enabling them to escape neutralizing antibodies in individuals immunized with the 1^st^ generation of spike protein based vaccines ^[26]^. Recent reports have emerged wherein N protein has been explored as a vaccine candidate and has displayed promising results. Immunization with N protein alone was shown to protect both mice and hamsters against SARS-CoV-2 viral challenge and a positive correlation was established between N peptide reactive T-cell response and the level of protection ^[27]^. Inclusion of N protein along with S protein for immunization has also been shown to aid viral clearance from distal organs like the brain compared to S protein alone ^[28]^. We observed that N protein is highly immunogenic and capable of inducing anti-N antibodies without addition of adjuvants. The splenocytes of RelCoVax® immunized animals showed a strong IFN-γ response to stimulation with N protein and RBD plus N protein compared to unstimulated controls. Stimulation with RBD alone did produce a slightly higher mean IFN-γ response but a high mouse to mouse variation was seen and overall it was not significant compared to unstimulated control. Here we have demonstrated the ability of N protein to strongly stimulate T cell responses with a Th1 bias. Other studies have also reported Th1 biased T-cell responses in NHP and mice models upon immunization with S protein or a modified RBD protein in combination with N protein with no evidence of vaccine associated enhanced respiratory disease (VAERD) ^[29,30]^.

With the emerging threat of new SARS-CoV-2 variants, a vaccine capable of providing broad protection against multiple strains is desired. We tested the ability of sera from immunized mice to neutralize two SARS-CoV-2 virus strains including the Delta variant of concern that is currently dominant across multiple regions around the globe. Efficient neutralization of both viral strains was observed in mice sera immunized with full dose of RelCoVax® comprising of 10 μg of each antigen with near identical PRNT50 titers indicating that the vaccine can protect efficiently against the more infectious Delta strain.

We have demonstrated that RelCoVax® is immunogenic and capable of stimulating both humoral and cellular immune responses in mice and potentially confer broader protection compared to only S protein targeted vaccines especially against emerging variants of SARS-CoV-2. Strong cellular immune responses stimulated by N protein have the potential to protect RelCoVax® immunized individuals in case of breakthrough infections through asymptomatic or mild manifestations of COVID-19. These data support further testing of RelCoVax® as a COVID-19 vaccine candidate in advanced preclinical and clinical settings.

## Acknowledgements

We would like to acknowledge Dr Ruta Kulkarni and her team at NIBEC, Pune for performing the pRNT50 tests with immunized mice sera provided by Reliance Life Sciences Pvt. Ltd.

## Author Contributions

VR conceptualized the vaccine and provided overall guidance, AP, GECVR, GK and KS designed the constructs, performed the cloning experiments and expression studies, GM, SP, NK, SSP, and VV developed the upstream and downstream processes for protein production as well as characterization, PK performed the SPR experiments, GECVR, AP, AG and GK performed formulation studies, SHR provided guidance for animal testing, RRL and KPR performed animal experiments, GECVR, AP, GK and AG developed and performed ELISA assays, PR provided technical guidance and designed the ELISpot experiments that were performed by PR, AP and AG. AP wrote the manuscript.

## Abbreviations

ACE2: angiotensin converting enzyme 2
CHO: Chinese hamster ovary
CI: confidence interval
COVID-19: Corona virus disease 19
CpG: cytosine phospho guanine
*E. coli*: *Escherichia coli*
ELISA: enzyme linked immunosorbent assay
ELISpot: Enzyme linked immunosorbent spot
GMT: geometric mean titer
IFN-γ: interferon-γ
IgG: immunoglobulin G
N: nucleocapsid
pfu: plaque forming units
pRNT: plaque reduction neutralization test
RBD: receptor binding domain
SARS-CoV-2: Severe acute respiratory syndrome Corona virus 2
S: spike

## Notes

### Competing Interest Statement

All authors are employees of Reliance Life Sciences Pvt. Ltd. and have no other competing financial interests or personal relationships to disclose.

## References

[1] Peiris JSM, Yuen KY, Osterhaus ADME, Stöhr K. The Severe Acute Respiratory Syndrome. N Engl J Med 2003;349:2431–41. https://doi.org/10.1056/NEJMra032498.

[2] de Groot RJ, Baker SC, Baric RS, et al. Middle East respiratory syndrome coronavirus (MERS-CoV): announcement of the Coronavirus Study Group. J Virol 2013;87:7790–2. https://doi.org/10.1128/JVI.01244-13.

[3] Wu Z, McGoogan JM. Characteristics of and Important Lessons From the Coronavirus Disease 2019 (COVID-19) Outbreak in China: Summary of a Report of 72 314 Cases From the Chinese Center for Disease Control and Prevention. JAMA 2020;323:1239–42. https://doi.org/10.1001/jama.2020.2648.

[4] Dong E, Du H, Gardner L. An interactive web-based dashboard to track COVID-19 in real time. Lancet Infect Dis 2020;20:533–4. https://doi.org/10.1016/s1473-3099(20)30120-1.

[5] Corey L, Mascola JR, Fauci AS, Collins FS. A strategic approach to COVID-19 vaccine R&D. Science 2020;368:948–50. https://doi.org/10.1126/science.abc5312.

[6] Kyriakidis NC, Lopez-Cortes A, Gonzalez EV, et al. SARS-CoV-2 vaccines strategies: a comprehensive review of phase 3 candidates. NPJ Vaccines 2021;6:28. https://doi.org/10.1038/s41541-021-00292-w.

[7] Lan J, Ge J, Yu J, et al. Structure of the SARS-CoV-2 spike receptor-binding domain bound to the ACE2 receptor. Nature 2020;581:215–20. https://doi.org/10.1038/s41586-020-2180-5.

[8] Shang J, Wan Y, Luo C, et al. Cell entry mechanisms of SARS-CoV-2. Proc Natl Acad Sci USA 2020; 117:11727–34. https://doi.org/10.1073/pnas.2003138117.

[9] Ravichandran S, Coyle EM, Klenow L, Tang J, Grubbs G, Liu S, Wang T, Golding H, Khurana S. Antibody signature induced by SARS-CoV-2 spike protein immunogens in rabbits. Sci Transl Med 2020;12:eabc3539. https://doi.org/10.1126/scitranslmed.abc3539.

[10] Premkumar L, Segovia-Chumbez B, Jadi R, Martinez DR, Raut R, Markmann A, Cornaby C, Bartelt L, Weiss S, Park Y, Edwards CE, Weimer E, Scherer EM, Rouphael N, Edupuganti S, et al. The receptor-binding domain of the viral spike protein is an immunodominant and highly specific target of antibodies in SARS-CoV-2 patients. Sci Immunol 2020;5:eabc8413. https://doi.org/10.1126/sciimmunol.abc8413.

[11] Garcia-Beltran WF, Lam EC, St Denis K, et al. Multiple SARS-CoV-2 variants escape neutralization by vaccine-induced humoral immunity. Cell 2021;184:2372–83. https://doi.org/10.1016/j.cell.2021.03.013.

[12] Mlcochova P, Kemp SA, Dhar MS, et al. SARS-CoV-2 B.1.617.2 Delta variant replication and immune evasion. Nature 2021;599:114–9. https://doi.org/10.1038/s41586-021-03944-y.

[13] Planas D, Veyer D, Baidaliuk A, et al. Reduced sensitivity of SARS-CoV-2 variant Delta to antibody neutralization. Nature 2021;596:276–80. https://doi.org/10.1038/s41586-021-03777-9.

[14] Dutta NK, Mazumdar K, Gordy JT. The Nucleocapsid Protein of SARS-CoV-2: a Target for Vaccine Development. J Virol 2020;94:e00647–20. https://doi.org/10.1128/JVI.00647-20.

[15] Kang S, Yang M, Hong Z, et al. Crystal structure of SARS-CoV-2 nucleocapsid protein RNA binding domain reveals potential unique drug targeting sites. Acta Pharm Sin B 2020;10:1228–38. https://doi.org/10.1016/j.apsb.2020.04.009.

[16] Le Bert N, Tan AT, Kunasegaran K, et al. SARS-CoV-2-specific T cell immunity in cases of COVID-19 and SARS, and uninfected controls. Nature 2020;584:457–62. https://doi.org/10.1038/s41586-020-2550-z.

[17] Art JF, Vander Straeten A, Dupont-Gillain CC. NaCl strongly modifies the physicochemical properties of aluminum hydroxide vaccine adjuvants. Int J Pharm 2016;517:226–33. https://doi.org/10.1016/j.ijpharm.2016.12.019.

[18] Gupta RK, Rost BE. Aluminum Compounds as Vaccine Adjuvants. In: O’Hagan DT, editor. Vaccine Adjuvants, Methods in Molecular Medicine, Totowa NJ: Springer; 2000, p. 65–89. https://doi.org/10.1385/1-59259-083-7:65.

[19] Campbell JD. Development of the CpG Adjuvant 1018: A Case Study. In: Fox C, editor. Vaccine Adjuvants, Methods in Molecular Biology, New York: Humana Press; 2017, p. 15–27. https://doi.org/10.1007/978-1-4939-6445-1_2.

[20] Wu F, Zhao S, Yu B, et al. A new coronavirus associated with human respiratory disease in China. Nature 2020;579:265–9. https://doi.org/10.1038/s41586-020-2008-3.

[21] Shang J, Ye G, Shi K, et al. Structural basis of receptor recognition by SARS-CoV-2. Nature 2020;581:221–4. https://doi.org/10.1038/s41586-020-2179-y.

[22] Barton MI, MacGowan SA, Kutuzov MA, Dushek O, Barton GJ, Van der Merwe PA. Effects of common mutations in the SARS-CoV-2 Spike RBD and its ligand, the human ACE2 receptor on binding affinity and kinetics. Elife 2021;10:e70658. https://doi.org/10.7554/eLife.70658.

[23] Chen WH, Du L, Chag SM, et al. Yeast-expressed recombinant protein of the receptor-binding domain in SARS-CoV spike protein with deglycosylated forms as a SARS vaccine candidate. Hum Vaccin Immunother 2014;10:648–58. https://doi.org/10.4161/hv.27464.

[24] Kuo TY, Lin MY, Coffman RL, et al. Development of CpG-adjuvanted stable prefusion SARS-CoV-2 spike antigen as a subunit vaccine against COVID-19. Sci Rep 2020;10:20085. https://doi.org/10.1038/s41598-020-77077-z.

[25] Liang JG, Su D, Song TZ, et al. S-Trimer, a COVID-19 subunit vaccine candidate, induces protective immunity in nonhuman primates. Nat Commun 2021;12:1346. https://doi.org/10.1038/s41467-021-21634-1.

[26] Tao K, Tzou PL, Nouhin J, et al. The biological and clinical significance of emerging SARS-CoV-2 variants. Nat Rev Genet 2021;22:757–73. https://doi.org/10.1038/s41576-021-00408-x.

[27] Matchett WE, Joag V, Stolley JM, et al. Cutting Edge: Nucleocapsid Vaccine Elicits Spike-Independent SARS-CoV-2 Protective Immunity. J Immunol 2021;207:376–9. https://doi.org/10.4049/jimmunol.2100421.

[28] Dangi T, Class J, Palacio N, et al. Combining spike- and nucleocapsid-based vaccines improves distal control of SARS-CoV-2. Cell Rep 2021;36:109664. https://doi.org/10.1016/j.celrep.2021.109664.

[29] Gabitzsch E, Safrit JT, Verma M, et al. Dual-Antigen COVID-19 Vaccine Subcutaneous Prime Delivery With Oral Boosts Protects NHP Against SARS-CoV-2 Challenge. Front Immunol 2021;12:729837. https://doi.org/10.3389/fimmu.2021.729837.

[30] Hong SH, Oh H, Park YW, et al. Immunization with RBD-P2 and N protects against SARS-CoV-2 in nonhuman primates. Sci Adv 2021;7. https://doi.org/10.1126/sciadv.abg7156.

